# Miniaturized Lysosome Enrichment through Selective Plasma Membrane Destabilization

**DOI:** 10.64898/2026.07.23.740280

**Authors:** Sofia Fajardo-Callejon, Dominic Winter

**Affiliations:** Oncological Proteometabolomics, Department Metabolism, Senescence and Autophagy, Research Center One Health Ruhr, University Hospital Essen, University of Duisburg-Essen, Virchowstrasse 171, 45147 Essen, Germany; Institute for Biochemistry and Molecular Biology, Medical Faculty, Rheinische Friedrich-Wilhelms-University of Bonn, Nussallee 11, 53115 Bonn, Germany

## Abstract

Organelle enrichment presents a prerequisite for the unbiased biochemical analysis of cellular subcellular compartments. The performance of this step is decisive for the experiment’s success, and despite the high sensitivity of downstream analytical strategies, such as enzymatic assays, western blotting, or mass spectrometry (MS)-based OMICs approaches, typically tens of millions of cells are required, standing in stark contrast. Here, we demonstrate that selective rupture of the plasma membrane constitutes a limiting factor for the reduction of cell numbers and present an approach to overcome this limitation through detergent-based plasma membrane destabilization, followed by mechanical homogenization. By combination of this strategy with two common methods for lysosome enrichment, namely superparamagnetic iron oxide nanoparticles (SPIONs) and immunoprecipitation via 3xHA-tagged TMEM192 (TMEM IP), we scale down lysosome enrichment to only half a million cells and demonstrate that reduced input cell numbers yield superior results with respect to sample purity and organelle proteome characterization.

**Highlights:**

- Release of intact lysosomes negatively correlates with cell concentration.
- Combination of detergent-based plasma membrane destabilization and mechanical homogenization increases lysosomal intactness from low cell numbers.
- Enrichment columns require a minimum sample input.
- Immunoprecipitation of intact lysosomes enables enrichment from lower cell numbers.
- Proteomics of low input lysosome enriched fractions identifies superior performance.

**Motivation:** Modern mass spectrometry (MS)-based proteomics strategies facilitate the detection and quantification of peptides and proteins with unprecedented sensitivity, enabling the analysis of low input samples down to single-cells. However, subcellular fractionation experiments typically require tens of millions of cells to achieve sufficient yield and purity, presenting a strong contrast. This restricts the application of organelle profiling to cell lines that can be grown in sufficient amounts, excluding many physiologically relevant species which are only available in small quantities. To be able to miniaturize subcellular fractionation experiments, it is of crucial importance to overcome this limitation and to miniaturize sample preparation strategies.

## Introduction

Recent advances in mass spectrometry (MS) instrumentation led to significant improvements for the detection of proteins, lipids, and metabolites, enabling the characterization of molecular features with unprecedented throughput and sensitivity^1^. While better ion handling and detection by MS instruments play a key role in this process, also innovations regarding sample preparation and liquid chromatography are decisive, as their automation, miniaturization, and adaptation to low input samples greatly reduce sample loss^1–3^. Combined, these innovations enable by now the comprehensive analyses of very small sample amounts, ranging from few to single cells with a performance which was unimaginable some years ago^4,5^. This presents a crucial development, as the performance of proteomics, lipidomics, and metabolomics analyses, unlike genomics/transcriptomics which benefit from oligonucleotide amplification, are constrained by the total abundance of the respective molecules, and hence the intrinsic sensitivity of the analytical setup^6^.

Current protocols for the preparation of subcellular and organelle-enriched fractions stand in stark contrast to these developments, as they typically require tens of millions of cells to achieve sufficient yield and purity for downstream analyses^7^. For example, typical inputs for differential centrifugation experiments are in the range of ∼20-50 million cells^7,8^, and for organelle-specific enrichment by immunoprecipitation (IP) or superparamagnetic iron oxide nanoparticles (SPIONs), typically ∼20 to 100 million cells per replicate are used^9–13^. This restricts organelle enrichment protocols to cells that can be obtained in sufficient amounts, such as immortalized cell lines or larger tissues. Consequentially, high-performance organelle proteomics workflows are typically not readily applicable to low abundant primary cells, those derived from induced pluripotent stem cells (iPSCs), or such sorted by fluorescence-activated/magnetic cell sorting (FACS/MACS)^14,15^. These cells, however, are often of crucial importance for the investigation of physiological processes and the characterization of disease-relevant mechanisms^14,15^. There is, therefore, an urgent need to scale down organelle enrichment experiments to leverage recent technological developments for the characterization of subcellular compartments in cells available in limited quantities.

An organelle receiving increased attention in recent years is the lysosome, whose perception in the scientific community is gradually transitioning from an unregulated “waste bag” to a control center of cellular metabolism^16,17^. Lysosomes are characterized by an acidic pH, which is maintained by the vacuolar H^+^ ATPase (v-ATPase) complex, and contain ∼60 hydrolases that break down a large variety of intra-/extracellular substrates delivered by the endo-/phagocytosis and autophagy pathways^18^. Importantly, lysosomes can degrade most types of biological structures, and their substrates range from single molecules to whole cells/organelles. The resulting building blocks are exported and recycled, which is essential for cellular homeostasis. In this context, the mechanistic target of rapamycin complex 1 (mTORC1), whose activity is regulated at the lysosomal surface in response to both cytosolic and lysosomal nutrient levels, is elementary to control the interplay between cellular anabolic and catabolic programs^17,19^. Given this crucial role lysosomes play for cellular homeostasis, it is not surprising that alterations of their function has been linked to a variety of diseases, including ∼70 rare inherited lysosomal storage disorders^20^, as well as common pathologies such as cancer and neurodegenerative diseases^21^.

For the MS-based proteomic, lipidomic, and metabolomic characterization of lysosomes, their enrichment proved to be of crucial importance, and two major approaches have been heavily utilized by the community in recent years: anti HA-IP from cells overexpressing the lysosomal membrane protein TMEM192-3xHA (TMEM IP) and SPIONs^22^. We showed previously that these methods outperform other lysosome enrichment strategies^10^ and various groups utilized them for the enrichment of lysosomes from a large variety of biological samples, including several cell lines (HEK293, HeLa, HuH-7, SH-S5Y5, MEF, NIH3T3)^23^, liver/brain tissue from mice^24,25^, and whole *C. elegans* animals^26^. As for all organelle enrichment experiments, also the enrichment of lysosomes requires large amounts of input material, restricting its use, and hence the investigation of lysosomal content, in low abundant cell types.

In the current study, we characterized and optimized the process of lysosome enrichment and identified a correlation of plasma membrane disruption efficiency with utilized cell numbers as decisive limiting factor for its miniaturization. We demonstrate that plasma membrane destabilization using the non-ionic detergent NP-40 allows for a strong reduction of numbers of required cells, and that low-input lysosome enrichment enables full lysosomal proteome coverage while reducing levels of contaminating proteins, therefore improving sample purity and data quality. The resulting protocol allows for the high-performance analysis of lysosomes from a fraction of the typically utilized input.

## Results

### Release of intact lysosomes negatively correlates with cell numbers

To enable for the enrichment of lysosomes (and other organelles), it is of crucial importance to selectively rupture the plasma membrane will preserving the membrane of the organelle(s) of interest. Lysosome enrichment experiments typically only perform satisfactory for high cell numbers, despite the very high sensitivity of downstream analytical procedures. We, therefore, hypothesized that plasma membrane disruption, and hence liberation of intact lysosomes, presents a limiting factor, and that the number of cells used as input plays a crucial role.

To investigate the relationship between cell numbers and lysosomal integrity, we enriched lysosomes from 24 and 6 million human embryonic kidney (HEK293) cells, equivalent to four and one confluent 10 cm tissue culture dishes, respectively, using SPIONs. In both cases, cells were homogenized following established procedures in 8 mL isolation buffer using a 15 mL glass dounce homogenizer, and enriched using magnetic force^10,22^ (Figure 1A). We assessed enrichment efficiency and lysosomal integrity using an enzymatic assay for the lysosomal luminal hydrolase β-hexosaminidase (HEXB) in the presence/absence of Triton X-100, reflecting total/ruptured lysosomes, respectively (Figure 1B, Table S1). Enrichment from 24 and 6 million HEK293 cells resulted in a lysosomal recovery rate of ∼42% and 19%, respectively (Figure 1C), reflecting the well-known reduction of performance for lower cell numbers. To further confirm that discrepancies were exclusively due to plasma membrane rupture and that intact lysosomes were enriched from the post-nuclear supernatant (PNS), we performed immunoblot analyses for markers of the Golgi apparatus, endoplasmic reticulum and mitochondria, as well as the lysosomal membrane (LIMP2) and lumen (CTSD, Figure 1D). While binding of unspecific proteins remained constant, we observed, in accordance with HEXB assays, that 6 million cells presented with reduced PNS abundance and hence enrichment efficiencies for both lysosomal membrane and luminal proteins. As SPIONs reside in the lysosomal lumen, rupture of the lysosomal membrane results in their release, and hence lower enrichment efficiency, confirming that reduced lysosome enrichment was due to organelle rupture during dounce homogenization. To further address this, we investigated, in addition to the PNS, also the total fraction (TF) including intact cells, with respect to contained intact/ruptured lysosomes by HEXB enzyme assays (Figure 1E and 1F). In these experiments, we observed a direct correlation between cell numbers and the presence of intact lysosomes in the PNS, further indicating that reduction of cell numbers results in increased rupture of both the plasma membrane and lysosomes. Additionally, they revealed that release of lysosomes from high cell numbers is sub-optimal, as the PNS did not contain the majority of lysosomes present in the TF, due to the presence of intact cells.

**Figure 1:**
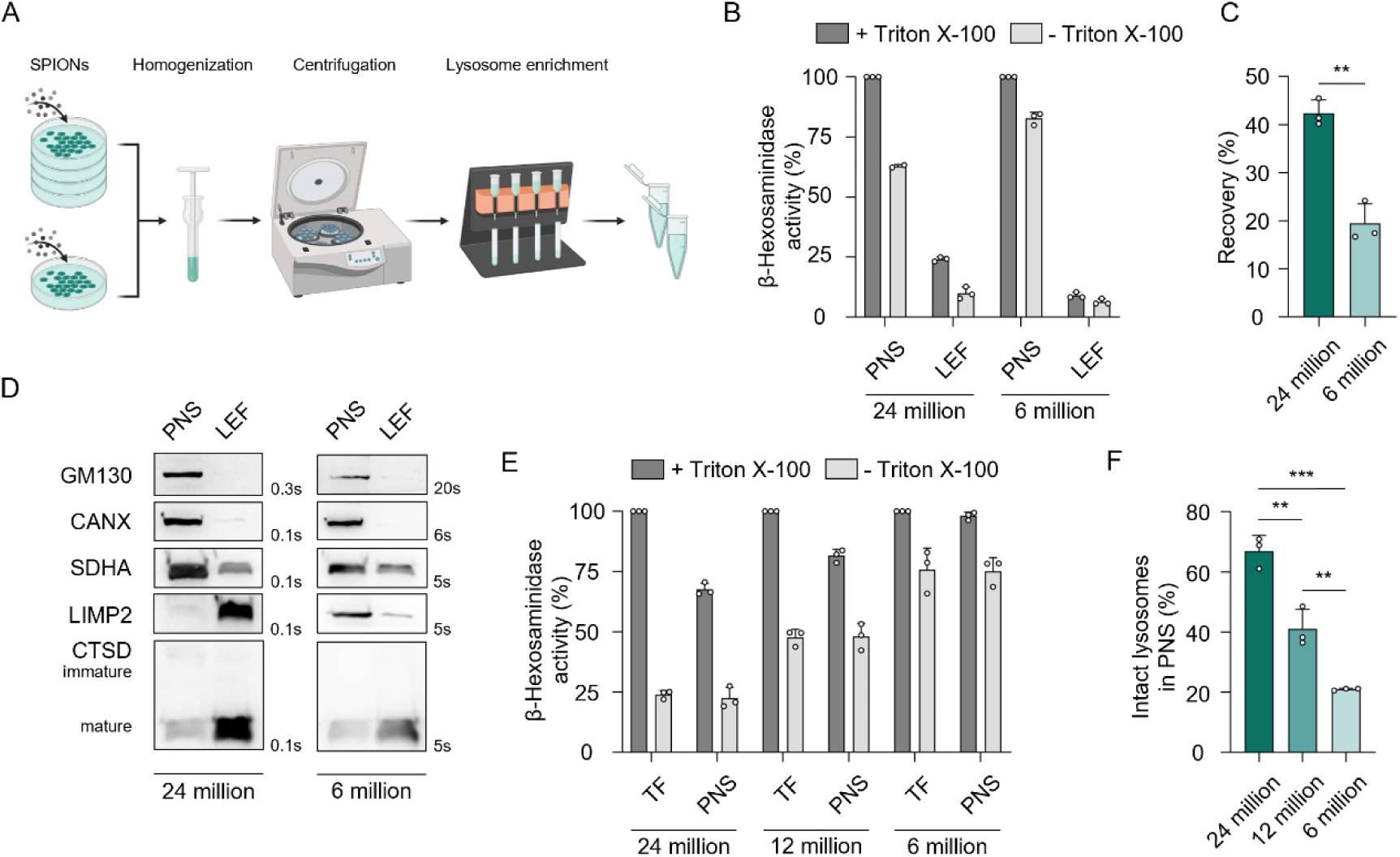
Cell numbers negatively correlate with enrichment of intact lysosomes. (A) Workflow for SPIONs lysosome enrichment from 24 and 6 million HEK293 cells. (B) Relative HEXB activity of individual SPIONs enrichment fractions. Addition of Triton X-100 enables assessment of lysosomal intact ratio. For raw values see Table S1. (C) Lysosomal recovery rates for 24/6 million cells. (D) Immunoblot analysis for marker proteins of lysosomes and contaminating organelles for lysosome enrichment experiments from 24 and 6 million HEK293 cells (Golgi apparatus: GM130, Endoplasmic reticulum: CANX, Mitochondria: SDHA, Lysosome: LIMP2, CTSD). (E) HEXB activity in TF/PNS for dounce homogenization of decreasing cell numbers. For raw values see Table S1. (F) Intact lysosomes present in the PNS of decreasing cell numbers. Shown are mean values ± SD, n = 3 (** = *p* < 0.01, *** = *p* < 0.001). TF = Total fraction, PNS = Post-nuclear supernatant, LEF = Lysosome-enriched fraction.

In summary, these experiments indicate that the number of cells used as input material for dounce homogenization is of crucial importance for the release of intact lysosomes, and hence their subsequent enrichment.

### Selective plasma membrane destabilization increases release of intact lysosomes

As homogenization of 24/12/6 million cells was performed in the same volume (8 mL), the cell density presented the only variable in these experiments. Thus, we reasoned that it could present the decisive factor for changes in the efficiency of selective plasma membrane rupture, as it was also pointed out previously as critical parameter^27^. To test this, we resuspended 6 million cells in either 8 mL or 1 mL and homogenized them using a 15 mL or 2 mL dounce homogenizer, respectively, followed by HEXB assays of individual fractions. Reduction of sample volume led to a 34% decrease of total enzymatic activity detected in the PNS compared to TF (Figure 2A), indicating that fewer cells were lysed; and a 2-fold increase of intact lysosomes (Figure 2B), confirming that cell density in the input fraction presents a critical factor. This finding explains the observed relationship and showcases a major problem of low input organelle enrichment, as further reduction of cell numbers would necessitate a concomitant reduction of sample volume, and hence dounce homogenizer dimensions, which presents with technical challenges.

**Figure 2:**
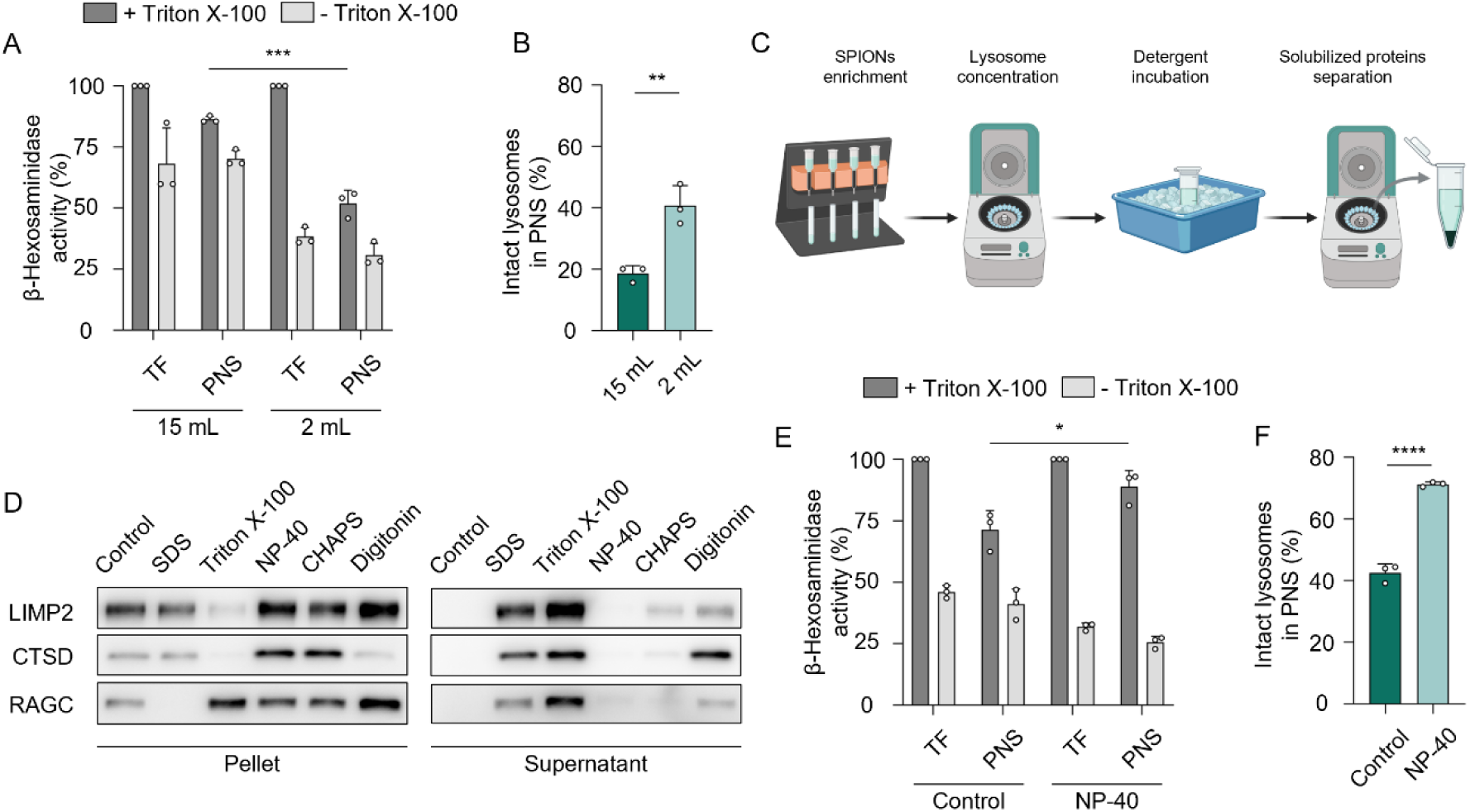
Improved release of lysosomes by homogenization volume reduction and plasma membrane destabilization. (A) TF/PNS HEXB activities for homogenization of 6 million HEK293 cells using a 15 mL or 2 mL glass dounce homogenizer. For raw values see Table S1. (B) Intact lysosome recovery in PNS fractions of different homogenization volumes. (C) Workflow for assessment of detergents with minimal effects towards lysosomal membrane permeabilization. (D) Immunoblot analysis for the effect of various detergents on lysosomal membrane integrity (lysosomal membrane: LIMP2, lumen: CTSD, associated: RAGC). (E) TNF/PNS HEXB activities for dounce homogenization of 6 million HEK293 cells with/without NP-40 pre-incubation. For raw values see Table S1. (F) Intact lysosomes in PNS of 6 million HEK293 cells with/without NP-40 pre-incubation. Shown are mean values ± SD, n = 3 (* = *p* < 0.05, ** = *p* < 0.01, *** = *p* < 0.001, **** = *p* < 0.0001). TF = Total fraction, PNS = Post-nuclear supernatant.

We, therefore, sought an alternative strategy to improve selective plasma membrane disruption irrespective of cell density of the input fraction. A common method in cell fractionation is the use of detergents, as their physicochemical properties enable disruption of hydrophobic/hydrophilic interactions^7,27^. Importantly, this allows for the selective weakening of distinct cellular membranes, as specific compartments typically present with unique lipid compositions. To identify a detergent that weakens the plasma membrane while preserving lysosomal intactness, we enriched lysosomes using SPIONs and incubated the resulting lysosome-enriched fractions (LEFs) with the commonly used detergents Sodium dodecyl sulfate (SDS), Triton X-100, Nonylphenol ethoxylate (NP-40), 3-[(3-Cholamidopropyl) dimethylammonio]-1-propanesulfonate (CHAPS), and Digitonin on ice. Subsequently, we precipitated lysosomes by centrifugation^10,28^, separating solubilized (supernatant) from intact (pellet) lysosomal membranes and used a luminal (CTSD), transmembrane (LIMP2), and associated (RAGC) lysosomal proteins as readout (Figure 2C). Only for samples treated with NP-40, all three proteins were present in the pellet and absent in the supernatant, indicating that it does not disrupt the lysosomal membrane, while for all other detergents various levels of membrane solubilization were observed. As NP-40 was utilized previously for the pre-treatment of intact cells to facilitate the enrichment of nuclei^29^, we concluded that treatment of cells with this detergent should enable destabilization of the plasma membrane while preserving lysosomal intactness.

We tested this by incubating 6 million HEK293 cells in a buffer containing 0.1% NP-40 for 30 min on ice before homogenization using a 2 mL dounce homogenizer. HEXB assays of the resulting fractions (Figure 2E) identified an increase in average lysosomal intact ratios from 42% to 71% (Figure 2F), surpassing even those obtained for 24 million cells (Figure 1E). Additionally, we observed a significant increase in PNS HEXB activity levels relative to the TF for cells incubated with NP-40 (Figure 2E), confirming more efficient plasma membrane disruption for combined NP-40 destabilization and dounce homogenization. Notably, higher NP-40 concentrations did not further increase lysosomal intact ratios (Figure S1A and S1B).

Together, these results demonstrate that reduction of sample volume and NP-40 incubation prior to dounce homogenization allow for selective plasma membrane disruption, and hence release of intact lysosomes, from smaller cell numbers

### NP-40-based plasma membrane destabilization facilitates lysosome enrichment from reduced cell numbers

We then benchmarked the performance of NP-40-assisted miniaturized lysosome enrichment using 6 million cells against the established procedure with 24 million cells using SPIONs. As expected, we obtained high lysosomal intact ratios in the PNS of 6 million cells (66%) using our optimized strategy, surpassing traditional homogenization of 24 million cells (45%), but lysosomal recovery and intactness was markedly lower (Figure 3A and Figure S2A).

**Figure 3:**
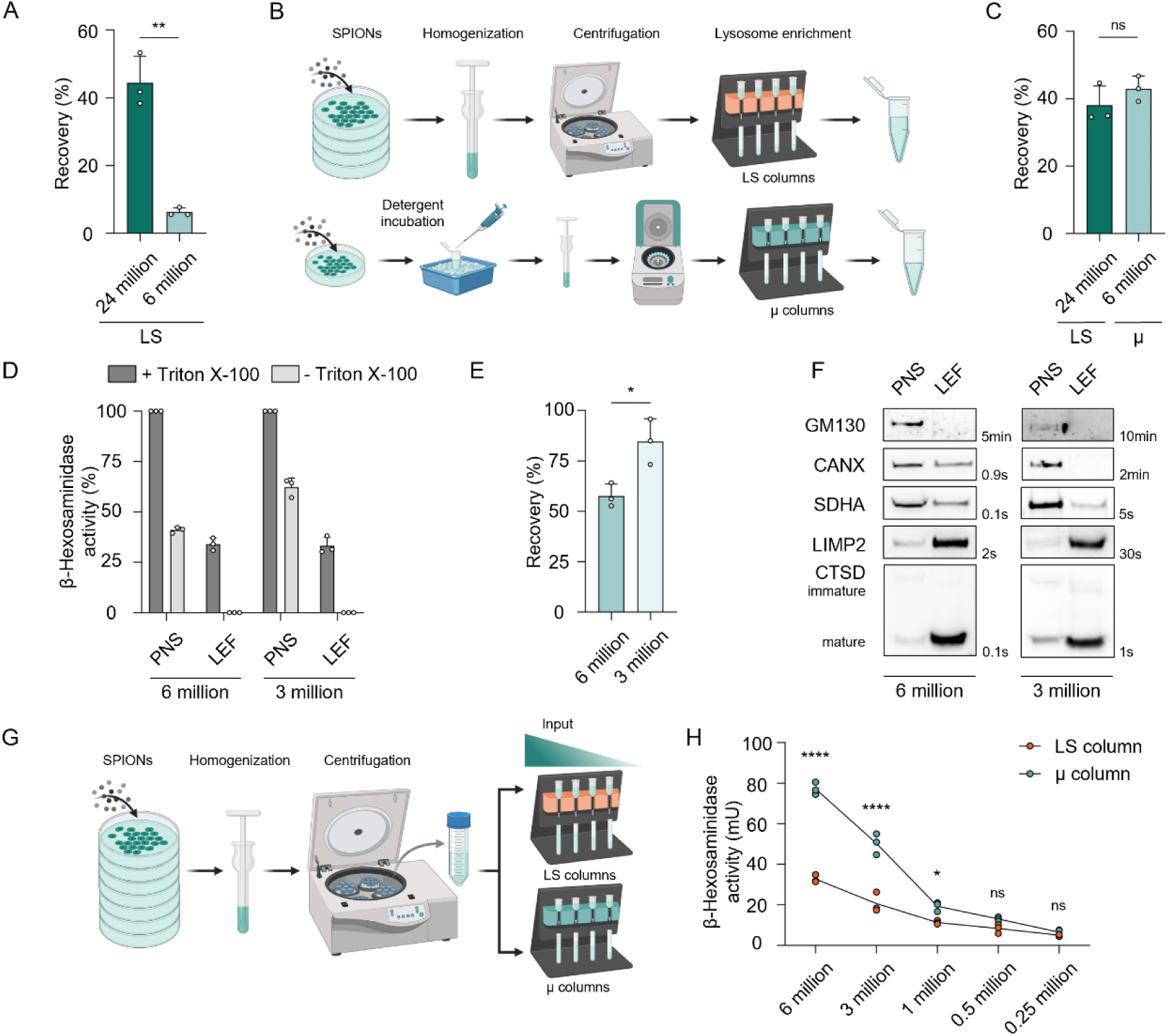
Plasma membrane destabilization enables lower input SPIONs enrichment of lysosomes. (A) HEK293 lysosomal recovery rates in LS column eluate fractions from 24 million cells dounce-homogenized using 15 mL and 6 million cells pre-treated with 0.1% NP-40 and dounce-homogenized using 2 mL. For raw values see Table S1. (B) Input-adjusted workflow for SPIONs enrichment from 24 million cells (LS columns) and 6 million cells (plasma membrane destabilization) and µ columns. (C) Lysosome recovery rates in eluate fractions obtained from 24 and 6 million cells using LS and µ columns, respectively. For raw values see Table S1. (D) HEXB activity in PNS/LEF for plasma membrane-destabilized and µ column-enriched lysosomes from 6 million and 3 million cells. For raw values see Table S1. (E) Lysosome recovery rates in eluate fractions from 6 million and 3 million cells using µ columns. (F) Immunoblot analysis of PNS/LEF from 6 million and 3 million HEK293 cells using miniaturized SPIONs enrichment (Golgi apparatus: GM130, Endoplasmic reticulum: CANX, Mitochondria: SDHA, Lysosome: LIMP2, CTSD). (G) Workflow for evaluation of enrichment column capacity. (H) HEXB activity in LEFs of SPIONs enrichment experiments with LS or µ columns of decreasing input amounts. For raw values see Table S1. Shown are mean values ± SD, n = 3 (* = *p* < 0.05, ** = *p* < 0.01, **** = *p* < 0.0001, ns = not significant). PNS = Post-nuclear supernatant, LEF = Lysosome-enriched fraction, LS = LS columns, µ = µ columns.

We demonstrated previously, that the utilized LS enrichment columns bind high amounts of unspecific proteins^23^. Therefore, we reasoned that the reduction of total input material could result in reduced saturation of plastic surfaces by other high-abundant non-lysosomal cellular components, leading to increased binding of lysosomes to unoccupied plastic surfaces, and hence their loss. Accordingly, we tested the utilization of a smaller column format (µ columns) designed for the enrichment of low input samples (Figure 3B). Due to their different design, these columns did not allow for the elution of intact lysosomes, because of which we eluted them with 1% Triton X-100, the standard procedure for TMEM IP experiments^25,30^. Combination of NP-40 incubation prior to homogenization with µ columns enabled the enrichment of lysosomes from 6 million cells with a recovery of 43%, comparable to values obtained for 24 million cells in combination with LS columns (Figure 3C and Figure S2B). Further reduction of cell numbers to 3 million, equivalent to one confluent 6 cm tissue culture dish, resulted in a lower lysosomal intact ratio compared to 6 million cells in the PNS (Figure 3D); however, average enrichment efficiencies increased, as most intact lysosomes present in the PNS were recovered (Figure 3E). Also, western blot analysis for marker proteins of contaminating organelles and lysosomes confirmed an excellent performance for the enrichment of lysosomes from 3 million cells (Figure 3F).

To validate that the observed effect is solely related to the usage of enrichment columns, and to evaluate the feasibility of lysosome enrichment from lower cell numbers, we further tested both column formats with fractions of one big batch of PNS to enable uncoupling of dounce homogenization and lysosome enrichment efficiencies. We, therefore, generated PNS fractions equivalent of 6/3/1/0.5/0.25 million cells from dounce homogenization of 24 million cells and enriched them using LS and µ columns, respectively (Figure 3G). These experiments confirmed the observed effect for 6 and 3 million cells, assigning the enrichment columns a decisive role, and demonstrated that further miniaturization with this setup is not feasible (Figure 3H).

Collectively, these results indicate that the combination of plasma membrane destabilization and µ columns enables SPIONs enrichment of lysosomes from one-eight of the starting material, and that 3 million cells present the lower limit of input for this strategy.

### Mass spectrometry-based proteomics reveals conserved organelle proteome composition for low input lysosome enriched fractions

To investigate the effect of reduced cell numbers on the composition of enriched lysosomes, we performed single-pot, solid-phase-enhanced sample preparation (SP3 digestion of LEFs from 24 and 3 million cells and analyzed them by data independent acquisition liquid chromatography tandem mass spectrometry (DIA-LC-MS/MS). In these experiments, we identified 6460 and 4659 proteins for LEFs obtained from 24 and 3 million cells, respectively, including 280 and 268 manually curated high-confidence lysosomal proteins^18^ (Figure 4A and 4B, Table S2). The reduction of total identified proteins by >1600 presented a strong contrast to the virtually similar numbers of such related to the lysosomes, and we interpreted these findings as further evidence for reduced binding of unspecific proteins to the plastic surface of enrichment columns (Figure 3). This, in turn, predicted that LEFs obtained from 3 million cells should present with reduced abundance of proteins originating from contaminating organelles relative to such located at lysosomes.

**Figure 4:**
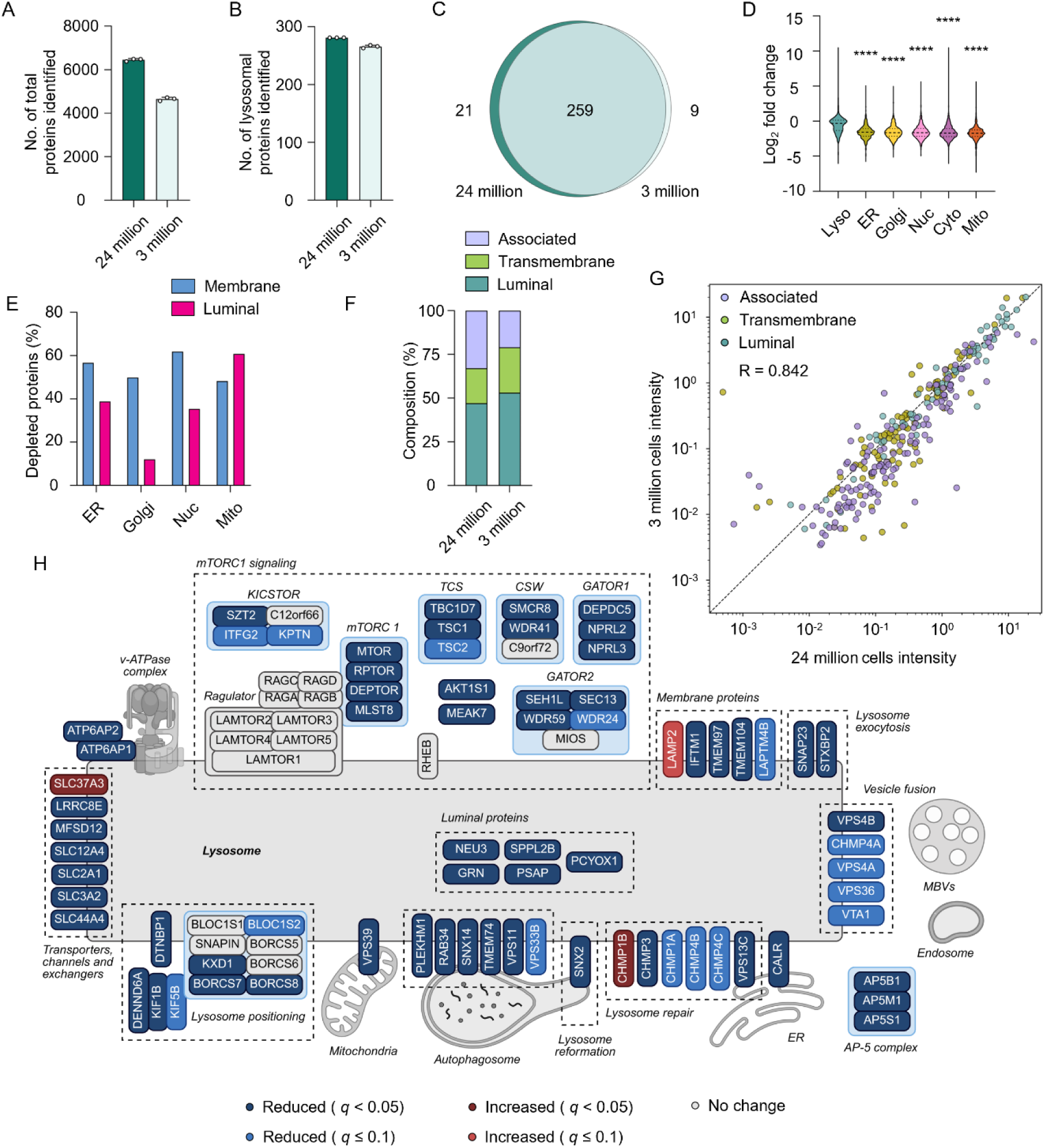
Miniaturization of SPIONs enrichment improves purity of lysosome-enriched fractions. (A/B) Identified total (A) and lysosomal (B) proteins for SPIONs-based lysosome enrichment from 24 million and 3 million cells. (C) Overlap of lysosomal proteins detected using 24 million or 3 million cells. (D) Distribution of log_2_ fold-change values for proteins assigned to distinct cellular compartments. (E) Percentage of depleted membrane and luminal proteins from different cellular compartments. (F) Sub-lysosomal protein composition dependent on localization. (G) Lysosomal protein abundance correlation for 24 million and 3 million cells (*R* = 0.842). (H) Differential abundance level overview of selected lysosomal proteins for 24 million and 3 million cells. Shown are mean values ± SD, n = 3 (**** = *p* < 0.0001, representing the comparison between each organelle and lysosomes). Mito = Mitochondria, Cyto = Cytoplasm, Nuc = Nucleus, Golgi = Golgi Apparatus, ER = Endoplasmic Reticulum, Lyso = Lysosome, MVBs = Multivesicular bodies.

To address this, we compared the abundances of lysosome-related proteins by replicate-wise normalization on mean v-ATPase complex subunit intensities^23^ (Table S2). The resulting contribution of individual organelles (based on GO classification and manually curated high-confidence lysosomal proteins^18^) revealed, in line with the notion of reduced unspecific binding, a decrease in signal intensity for proteins from all organelles except the lysosome, with highest depletion factors for the nucleus and cytosol (Figure 4D and Figure S3A to S3F, Table S3). We further assessed the sub-organellar dynamics of contaminants to assess if rather intact organelles, or fractions thereof, were depleted (Table S4). Interestingly, we observed a stronger depletion of membrane proteins relative to such located in the respective lumen of the nucleus, Golgi, and ER, reminiscent of an unspecific enrichment of membrane fragments rather than of intact organelles or the effect of Triton X-100 used for lysosome enrichment form µ columns. For mitochondria, on the other hand, we found similar values for different sub-organellar compartments, with slightly higher depletion factors for matrix proteins, indicating that mainly intact organelles were unspecifically enriched (Figure 4E). Cytosolic proteins presented generally with rather low fold-change values compared to organellar proteins, demonstrating that their interaction with plastic surfaces does not seem to present a major contaminant, probably due to their hydrophilic nature, and that probably many of the detected proteins specifically interact with lysosomes.

Further characterization of lysosomes with respect to the sub-organellar localization of their proteins identified a presumable change in composition between LEFs from 24 and 3 million cells, mainly due to the reduced abundance of proteins dynamically interacting with lysosomes (lysosome associated proteins), as well as few located in the lysosomal lumen and membrane^21^ (Figure 4F and 4G, Figure S3G to S3I)). Detailed manual investigation of these proteins identified almost exclusively downregulation of lysosomal membrane and luminal proteins previously reported with dual localization, i.e. such which are not only present at lysosomes but also at other subcellular compartments^31^, as well as lysosome-associated complexes which dynamically interact with lysosomes, but are predominantly cytosolic (Figure 4H). For instance, proteins related to autophagosomes (TMEM74^32^), lysosomal repair (CHMP1A, CHMP3, CHMP4A/B/C, VPS13C^33,34^), formation of contact sites with other organelles (VPS39^35^), as well as such involved in cargo delivery and vesicle-fusion (PLEKHM1, RAB34, SNX14, VPS11, VPS4B^36–38^) were generally of lower abundance in LEFs from 3 million cells (Figure 4G, Table S3). In accordance, for mTORC1, we observed member-specific dynamics based on their residence at the lysosome. The cytosolic mTORC1 components KICSTOR, TSC1/2, and GATOR1/2 were of lower abundance, while the Rag GTPases^39,40^, which are tightly associated with the membrane-bound Ragulator complex^41,42^, showed no significant changes. Curiously, also for the lysosomal complexes v-ATPase and BLOC-one-related complex (BORC), which are typically only discussed with respect to lysosomes as fully assembled (sub-) complexes, we observed lower abundance for certain subunits. For the v-ATPase, the accessory subunits ATP6AP1 and ATP6AP2 presented with decreased abundance. While both of them have been linked to lysosomal biogenesis and assembly of the v-ATPase complex prior to its release from the *trans*-Golgi network^43^, ATP6AP2 was further shown to participate in Wnt/β-catenin signalling at the plasma membrane^44^, providing a likely explanation for their reduced abundance at lysosomes. Similarly, three out of eight BORC subunits (KXD1, BORCS7 and BORCS8) were reduced in abundance, while the other subunits, together with ARL8A/B, small GTPases recruited to lysosomes by BORC, remained constant. In line with these findings, a recent study described hierarchical assembly of BORC, where subunits BLOC1S1, BLOC1S2, SNAPIN and BORCS5 forms a core scaffold that recruits the other subunits^45^, indicating that they are present in excess in the cytosol. Collectively, these results demonstrate that the miniaturized SPIONs protocol enables the enrichment of lysosomes from a fraction of the starting material, with comprehensive coverage of their core proteome at reduced levels of proteins originating from contaminating compartments.

### Miniaturized TMEM IP enables lysosome enrichment from low cell numbers

As the utilized columns’ properties prevented further miniaturization of SPIONs-based lysosome enrichment (Figure 3H), we argued that TMEM IP^10,25,30,46,47^, which is based on the binding of lysosomes to magnetic immunoaffinity beads, could allow for further downscaling of the process. We first tested if it is possible to dounce-homogenize <3 million cells, as this presented a key requirement. We were able to obtain satisfactory lysosome intact ratios (∼30%) also in the PNS of 1 and 0.5 million cells (albeit at lower rates then for 3 million cells, Figure 5A and 5B). As the obtained amounts of liberated intact lysosomes, however, were sufficient for comprehensive downstream analyses, we proceeded with lysosome enrichment from 0.5 million cells by TMEM IP (Figure 5C and 5D). Strikingly, we were able to recover 84% of intact lysosomes from 0.5 million cells, compared to 68% for 18 million cells, the input typically utilized for TMEM IPs^46^ (Figure 5E). Western blot analyses of LEFs from these experiments showed not only the enrichment of lysosomal markers, but also a concomitant reduction of proteins originating from contaminating cellular compartments (Figure 5F), indicating that miniaturized TMEM IP generates higher purity LEFs than the typically utilized high-input experiments, analogous to the results obtained from SPIONs (Figure 4).

**Figure 5:**
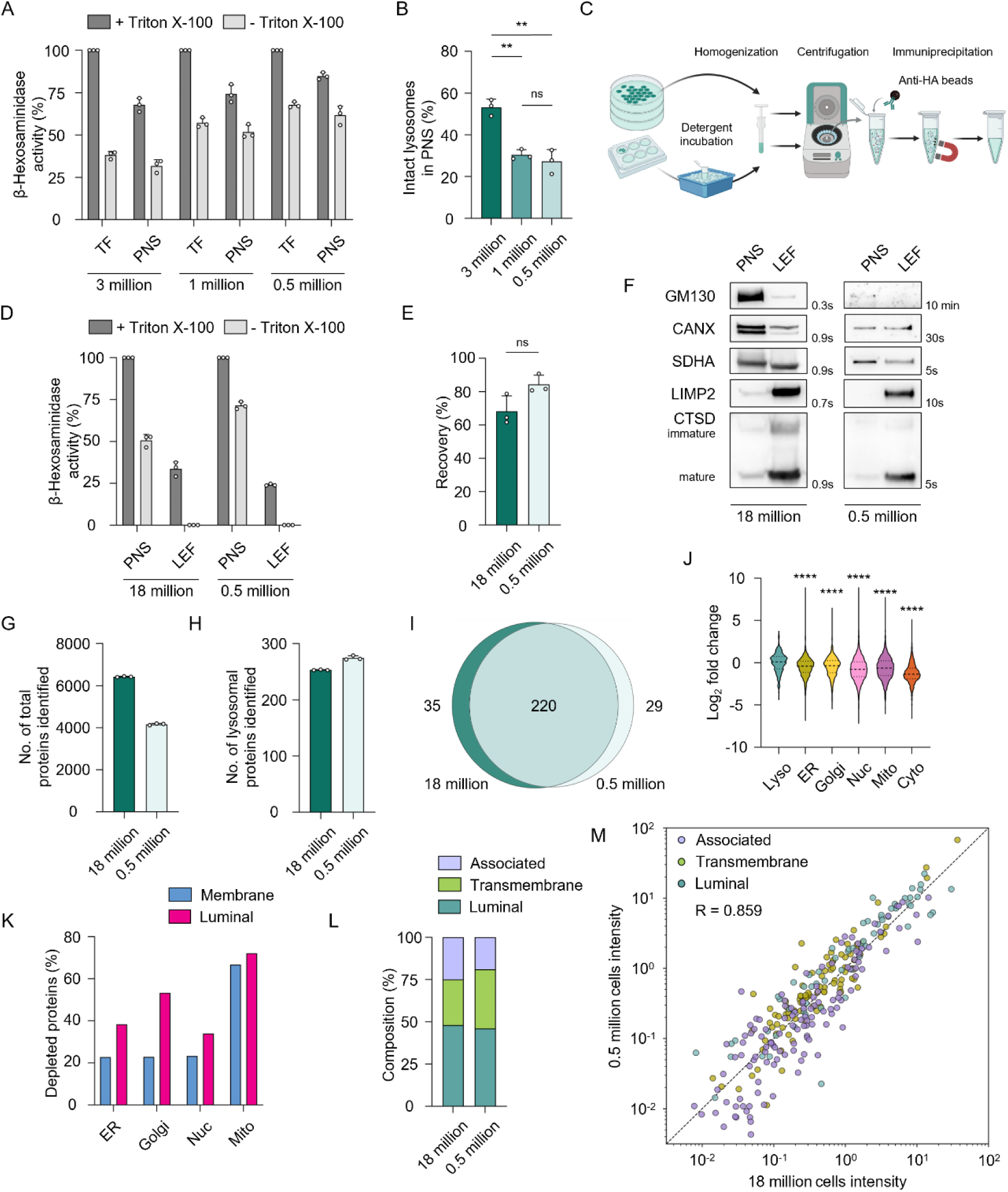
Miniaturized TMEM IP enables enrichment of lysosomes with increased purity from low cell numbers. (A/B) TF/PNS HEXB activity (A) and lysosomal intactness (B) for dounce homogenization after NP-40 incubation of 3/1/0.5 million HEK293 TMEM cells. For raw values see Table S1. (C) Workflow for lysosome enrichment by TMEM IP from HEK293 TMEM cells with/without plasma membrane destabilization. (D/E) HEXB activity (D) and lysosome recovery rate (E) for lysosome enrichment experiments from 18 million and 0.5 million HEK293 TMEM cells. For raw values see Table S1. (F) Immunoblot analysis of PNS/LEFs from 18 million and 0.5 million HEK293 TMEM cells (Golgi apparatus: GM130, Endoplasmic reticulum: CANX, Mitochondria: SDHA, Lysosome: LIMP2, CTSD). (G/H) Identified total (G) and lysosomal (H) proteins for TMEM IP lysosome enrichment from 18 million and 0.5 million cells. (I) Overlap of lysosomal proteins detected using 18 million or 0.5 million cells. (J) Distribution of log_2_ fold-change values for proteins assigned to distinct cellular compartments. (K) Percentage of depletion of membrane and luminal proteins from different cellular compartments. (L) Sub-lysosomal protein composition dependent on localization. (M) Lysosomal protein abundance correlation for 18 million and 0.5 million cells (*R* = 0.842). Shown are mean values ± SD, n = 3 (** = *p* < 0.01, **** = *p* < 0.0001, representing the comparison between each organelle and lysosomes, ns = not significant). TF = Total fraction, PNS = Post-nuclear supernatant, LEF = Lysosome-enriched fraction. Mito = Mitochondria, Cyto = Cytoplasm, Nuc = Nucleus, Golgi = Golgi Apparatus, ER = Endoplasmic Reticulum, Lyso = Lysosome

To further follow up on this finding, we performed SP3 digestion of 10 µg LEF protein from 18 or 0.5 million cells and analyzed the samples by DIA-LC-MS/MS, identifying 6527 and 4197 proteins respectively (Figure 5G, Table S5). Notably, 278 lysosomal proteins were identified using 0.5 million cells, compared to 255 detected from 18 million (Figure 5H-I, Table S5). Furthermore, lysosome-centric v-ATPase-normalized fold-change values revealed increased intensities for lysosomal proteins and a markedly reduced abundance of proteins localized at other subcellular compartments, with mitochondria, nucleus and cytosol showing highest depletion factors (Figure 5J, Table S5 and Table S6). This indicates, analogous to SPIONs (Figure 4), that miniaturization of lysosome enrichment by TMEM IP increases the purity of enriched lysosomes. In contrast to SPIONs miniaturization, we observed stronger depletion of contaminating organelles’ luminal proteins, probably because elution of lysosomes from affinity beads was performed in both instances with Triton X-100 and therefore the interaction of solubilized membrane proteins with affinity beads was similar (Figure 5K). Depletion of non-lysosomal intracellular vesicles was in general higher than for SPIONs miniaturization, while proteins from the endolysosomal compartment were virtually constant (Table S7), in line with previous results from our group demonstrating increased enrichment of non-lysosomal vesicular fractions by TMEM IPs^22^. Further comparison of individual protein dynamics based on their subcellular localizations (Figure S5) identified lower reduction of background proteins relative to SPIONs experiments (Figure S3) but also increased enrichment of such located at the lysosome. Regarding distinct lysosomal protein populations, we found that all lysosomal proteins showed a similar behavior. This matched those of luminal and associated proteins for SPIONs miniaturization, but presented with increased recovery of membrane proteins (Figure 5L and 5M, Figure S5 G-I). A possible explanation for this observation is that similar conditions are used for miniaturized TMEM IPs, while different column formats are utilized for SPIONs, resulting in more reproducible conditions. This demonstrates that miniaturized TMEM IP enables the enrichment of lysosomes with increased coverage and purity and no loss of performance.

Taken together, both SPIONs and TMEM IP experiments demonstrate that rupture of the plasma membrane presents the limiting factor for further downscaling of lysosome enrichment. Combination of detergent-based membrane destabilization with mechanical disruption, presents an efficient strategy to overcome this limitation. Using miniaturized lysosome enrichment, we were able to enrich lysosomes from 500,000 cells presenting a 48-fold reduction of starting material relative to currently widely used protocols.

## Discussion

Here, we present a strategy that combines detergent-based pre-treatment of cells with mechanical homogenization, to efficiently disrupt the plasma membrane while maintaining lysosomal integrity. This enables the enrichment of lysosomes from a fraction of cell numbers typically utilized as input, as little as half a million cells, presenting a marked decrease of material compared to other widely used approaches.

We identified the mechanical homogenization of cells, for which we used a glass dounce homogenizer, as a key step for the miniaturization of lysosome enrichment. The size of this device was of crucial importance, implying that smaller dounce homogenizers or other devices, such as ball-bearing homogenizers^48^ could enable further miniaturization. Another important factor was the choice of detergent, with NP-40 yielding ideal results. NP-40 is known as a mild detergent, widely conserving native protein conformation^27^, and has been used previously in low concentrations for the isolation of nuclei, as it specifically solubilized other organelles, such as the ER, Golgi apparatus and mitochondria^29^. A likely reason for the excellent performance for nuclei enrichment in these experiments was the specific composition of the largely detergent-resistant nuclear lamina^49^, which is probably also the case for our results regarding the lysosomal membrane, as the latter has a distinct lipid composition and is rich in highly glycosylated lysosome-associated membrane proteins (LAMPs)^50^. The sugar moieties of these proteins’ luminal domains form the lysosomal luminal glycocalyx, which protects the lysosome from self-digestion^18,51,52^ but also provides mechanical stability. This could be decisive in the context of stability towards NP-40 in low concentrations.

It is, however, also possible to dissolve the lysosomal membrane with NP-40, as it has been recently used for the elution of samples after affinity enrichment of lysosomes in RPE-1 cells^53^. This showcases that the choice of the detergent type and concentration is crucially dependent on the cell line utilized, and it is highly likely that adaptation of miniaturized lysosome enrichment to other types of cells could benefit from testing different detergents in various concentrations to yield optimal results. Importantly, as we used NP-40 only for pre-treatment of intact cells, and samples were prepared for MS analysis via SP3, we did not observe any adverse effects on sample purity for downstream proteomics workflows from its usage, presenting a high compatibility with OMICs workflows.

In our DIA-MS analyses, we identified that miniaturization of lysosome enrichment resulted in higher LEF purity while preserving (SPIONs) or increasing (TMEM IP) the numbers of identified and quantified lysosomal proteins relative to conventional sample preparation. While this confirms that the limitation for the miniaturization for lysosome enrichment solely depends on the selective rupture of the plasma membrane, it also showcases that reduction of cell numbers for lysosome enrichment improves the performance of downstream MS analyses, rather than reducing it, due to reduction of input material. This is most likely due to reduced unspecific binding of unrelated proteins and organelles to plastic surfaces. Furthermore, for TMEM IPs, this reduction in complexity probably enabled the identification of more lysosomal proteins, as we demonstrated before that overall sample complexity presents a major limiting factor for the identification of low abundant lysosomal proteins^10,54,55^.

Strikingly, we found in these analyses for both SPIONs and TMEM IPs not only a reduction in intensity for non-lysosomal contaminating proteins but also for lysosome-associated ones. This could either be the result of predominantly cytosolic localization of these proteins, indicating that large portions of them actually can be seen as background (despite their partial lysosomal localization), or a lack of lysosomal recruitment, as consequence of enrichment-specific modulation of lysosomal properties. As it was shown in various previous studies that fully functional lysosomes are recovered by both SPIONs and TMEM IP^56–58^, the latter presents an unlikely scenario, indicating that LEFs obtained from smaller amounts of input present a more realistic picture of lysosomal composition compared to large cell numbers.

Furthermore, the lysosomal abundance of fully lysosome-localized luminal and transmembrane proteins was not affected, demonstrating recovery of intact organelles which is in line with our HEXB assay results. We did observe, however, reduced abundance of lysosomal proteins that we recently identified to present with a dual localization, as part of the abovementioned reduction of other contaminating organelles, further confirming that they do not exclusively reside at the lysosome^31^. This demonstrates that our miniaturized protocols can consistently enrich and detect lysosomal luminal proteins and such embedded- or closely associated with the lysosomal membrane. Importantly, they are likely to present with a better performance to detect abundance changes at the lysosomal membrane, as the influence of differences in expression affecting the total protein pool (including other non-lysosomal localizations) are less prominent.

Regarding differences in SPIONs and TMEM IP miniaturization, we observed more pronounced general downregulation of proteins for SPIONs experiments, which we currently attribute to the bigger differences in plastic surface areas between enrichment columns and the usage of Triton X-100 for lysosome elution from µ columns. Importantly, this also affected lysosomal proteins, in line with the notion that not only contaminating organelles but also lysosomes were unspecifically bound to said surfaces. In accordance, we failed to efficiently downscale lysosome enrichment using LS columns, which present a rather large surface area, and only the use of µ columns yielded satisfactory results. This limitation, which is irrespective of homogenization efficiency, restricts the further miniaturization of lysosome enrichment to IP-based approaches, which we demonstrated for TMEM IPs, a frequently used method for the enrichment of lysosomes^10,57^, based on the overexpression of 3xHA-tagged TMEM192.

As a previous study of our group demonstrated recently that TMEM192 overexpression significantly alters lysosomal composition in HEK293 cells^22^, endogenous Lyso IP^59,60^, or knock in cell lines, which both do not require overexpression of lysosomal membrane proteins^61^, present attractive alternatives for future studies. Combining lysosome enrichment miniaturization using NP-40-based plasma membrane destabilization with endogenous Lyso IP could, hence, pave the way for the investigation of more elusive samples, such as acutely isolated cells or primary cells grown in culture. These cells neither permit growth in large quantities, nor the generation of versions stably overexpressing tagged proteins, making them largely inaccessible to the currently widely used protocols. In accordance there is, to our knowledge, currently only one study describing the isolation of lysosomes from neuronal cultures^62^. Our strategy for the detergent-based selective destabilization of the plasma membrane could provide a viable option in this context, and optimization of individual detergent types and concentrations promises to enable its use not only in HEK293 cells, as demonstrated here, but also various other systems. As our approach does not add time or cost constraints to experimental procedures, it should present an attractive option for such experiments.

## Limitations of the study

This study was performed in HEK293 cells, which typically enable high organelle enrichment efficiencies. It is important to note, that other cells could require different input numbers to obtain a satisfactory performance, and that it is highly likely that protocol adaptation to individual cell types is required. This also applies for mechanical homogenization. While, in our hands, a glass dounce homogenizer presented a good solution, it is difficult to speculate if other homogenization techniques may benefit from miniaturized enrichment and what is most suitable for distinct cell types. Individual adaptation of protocols will be most likely necessary.

## Acknowledgments

We thank Selin Özesen Kozok, Dhriti Arora and Octavio Reyes-Matte for valuable discussions. This study was supported by the German Academic Exchange Service (DAAD) and the Heisenberg Program of the German Research Council (DFG). Figures were created with BioRender.com.

## Author contributions

**Sofía Fajardo-Callejón:** Conceptualization, Methodology, Software, Validation, Formal analysis, Investigation, Data curation, Writing – Original draft, Writing – Review & editing, Visualization, Project administration, Funding acquisition. **Dominic Winter:** Conceptualization, Methodology, Validation, Formal analysis, Investigation, Resources, Data curation, Writing – Original draft, Writing – Review & editing, Supervision, Project administration, Funding acquisition.

## METHODS

Detailed methods are provided in the online version of this paper and include the following:

- RESOURCE AVAILABILITY

`o` Lead contact
`o` Materials availability
`o` Data and code availability KEY RESOURCE TABLE
- EXPERIMENTAL MODEL AND SUBJECT DETAILS

`o` Cell lines
- METHOD DETAILS

`o` Enrichment of Lysosomes with Superparamagnetic Iron Oxide Nanoparticles (SPIONs)
`o` Enrichment of Lysosomes by TMEM192-3xHA immunoprecipitation (TMEM IP)
`o` β-Hexosaminidase Assay
`o` Immunoblotting
`o` Sample Preparation for Mass Spectrometry
`o` Liquid Chromatography-Tandem Mass Spectrometry Analysis
- QUANTIFICATION AND STATISTICAL ANALYSIS

`o` Mass Spectrometry Data Analysis
`o` Data Processing and Statistical Analysis of β-Hexosaminidase Assay

## RESOURCE AVAILABILITY

### Lead contact

Further information for resources and request should be directed to and will be fulfilled by the lead contact, Dominic Winter

### Materials availability

All the unique reagents generated in this study are available from the lead contact without restriction.

### Data and code availability

- The mass spectrometry proteomics data have been deposited to the ProteomeXchange Consortium *via* the PRIDE^63^ partner repository and are available with the dataset identified PXD081210.
- This study does not report original code.
- Any additional information required to reanalyze the data reported in this study is available from the lead contact upon request.

## STAR METHODS

### KEY RESOURCE TABLE

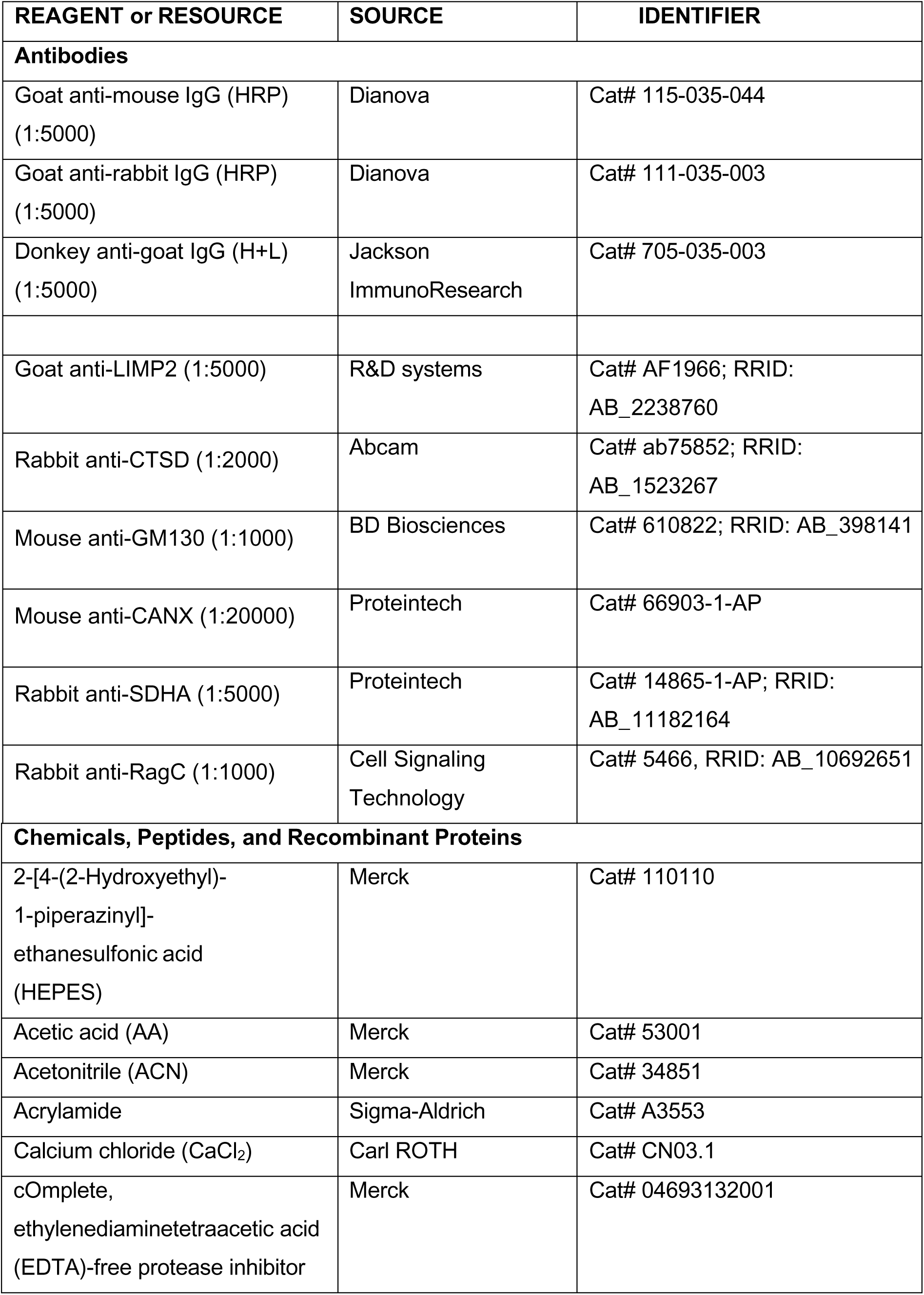
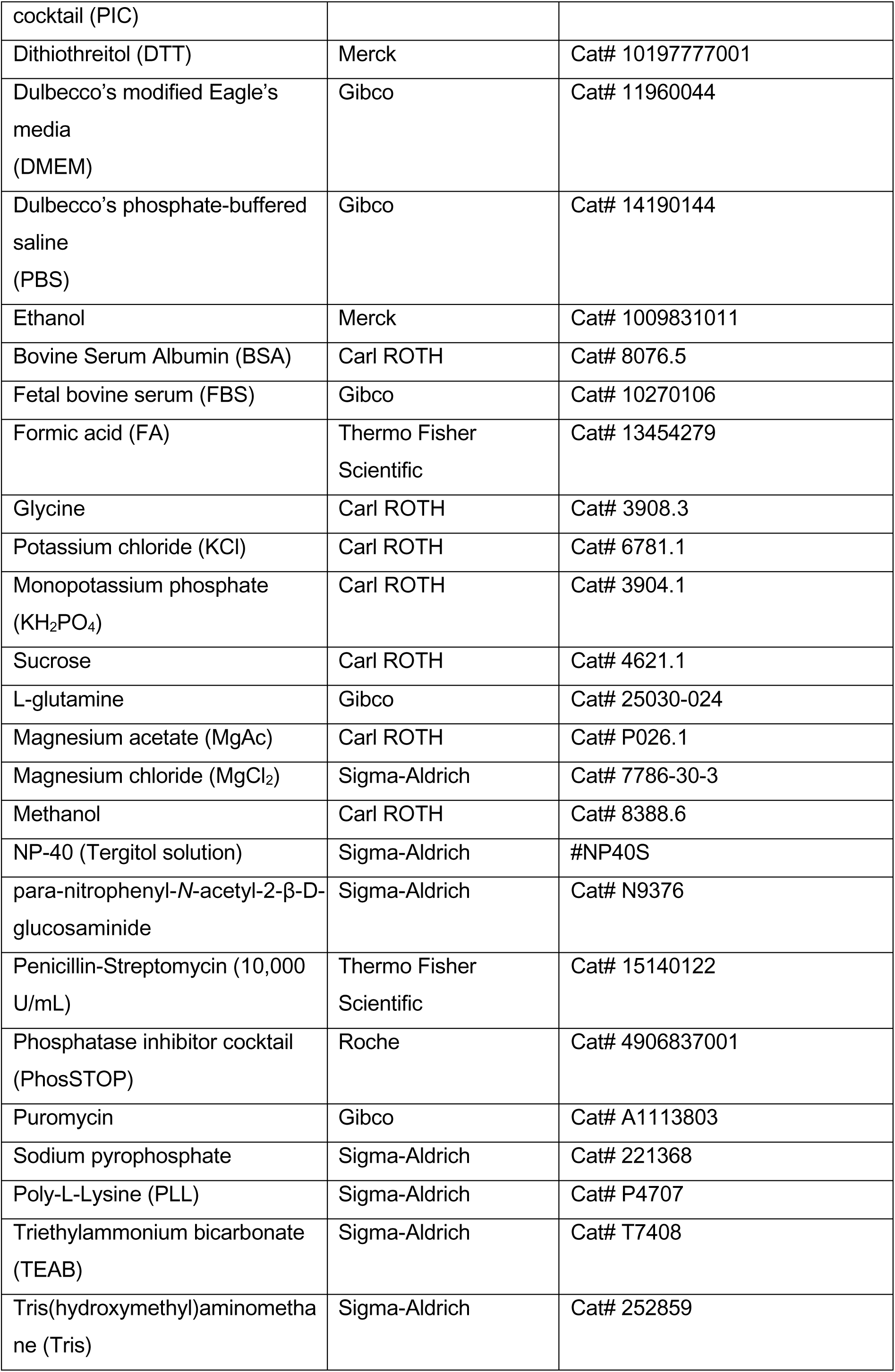
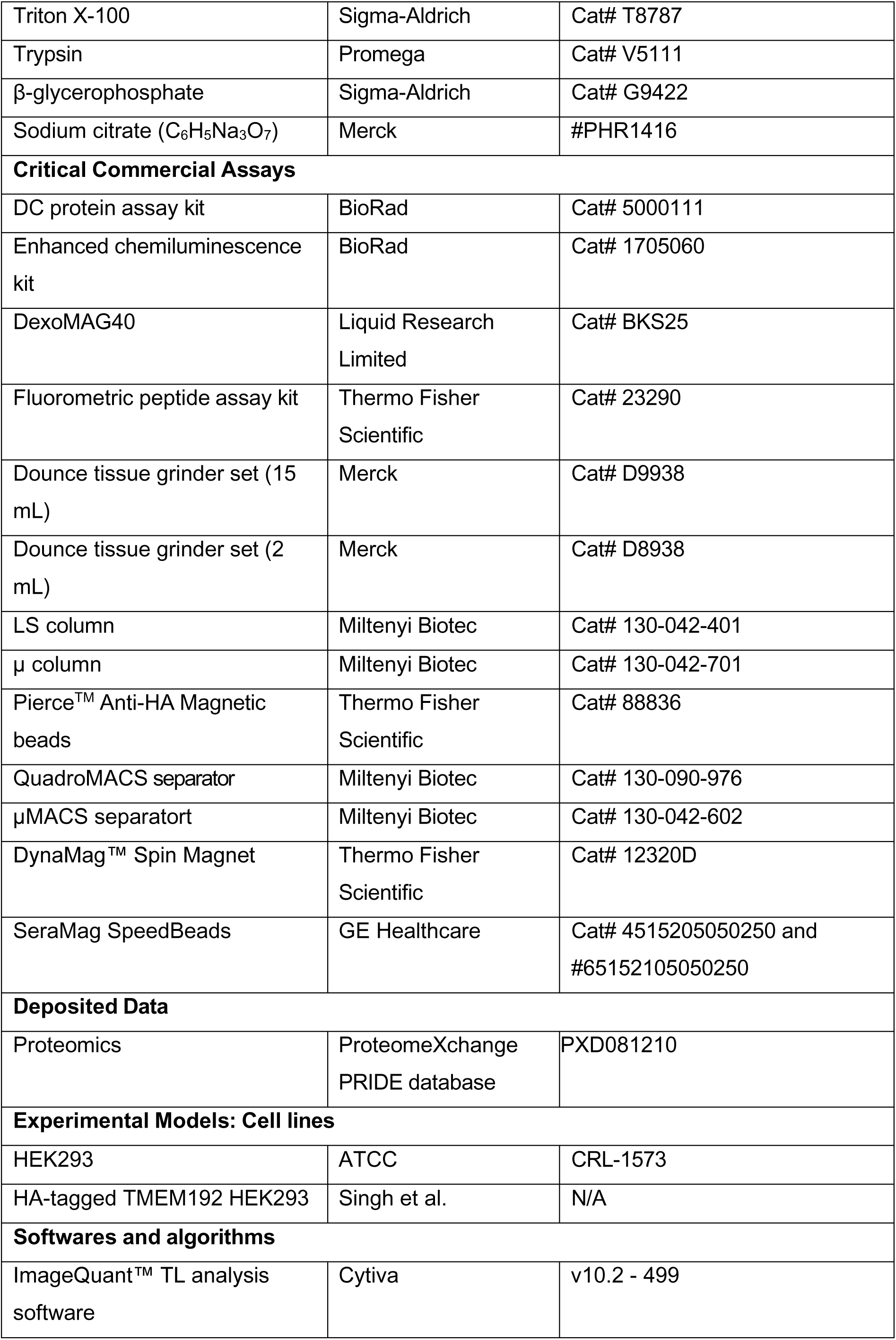
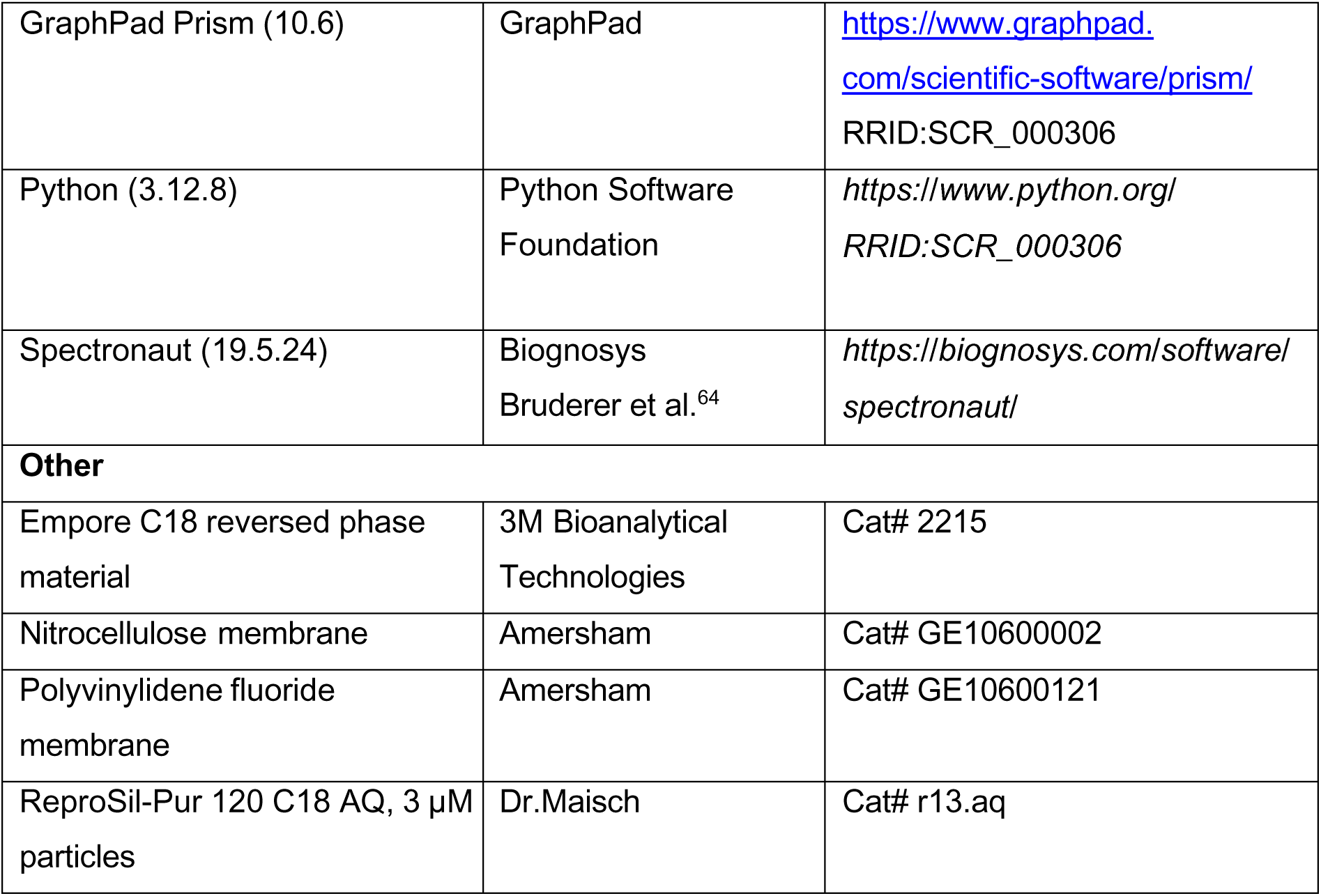

## EXPERIMENTAL MODEL AND SUBJECT DETAILS

### Cell lines

HEK293 WT and HEK293 cells expressing 3xHA-tagged TMEM192 (HEK293 TMEM) were cultured in Dulbecco’s modified Eagle’s media (DMEM) supplemented with 10% Fetal bovine serum (FBS), 100 µg/mL penicillin, 100 µg/mL streptomycin, 2 mM L-glutamine (complete medium) at 37 °C, and 5% CO_2_. All cell lines were regularly controlled for mycoplasma contamination using polymerase chain reaction (PCR).

## METHOD DETAILS

Refer to the key resources table for antibodies, chemicals, and software.

### Enrichment of Lysosomes with Superparamagnetic Iron Oxide Nanoparticles (SPIONs)

For lysosome enrichment, 10 cm tissue culture dishes were coated with 0.5 mg/mL Poly-L-Lysine (PLL) in 1X phosphate-buffered saline (PBS) for 25 min at 37 °C. HEK293 WT cells were seeded in complete medium supplemented with 10% SPIONs (Dextran: 40 kDa, core size: 8 nm, hydrodynamic size: 50 nm, Fe content: 10 mg/mL, solvent: dH_2_O). After a pulse period of 24 h, the medium was removed, cells were washed with 1X PBS, fresh complete medium was added, and cells were incubated for a chase period of 24 h. Subsequently, cells were washed twice with ice-cold 1X PBS, scraped in 8 mL of isolation buffer (250 mM sucrose, 10 mM HEPES/NaOH pH 7.4, 15 mM KCl, 1 mM MgCl_2_, 1.5 mM MgAc, 1X PIC), and lysed using a 15 mL dounce homogenizer (tight pestle) using 25 strokes. Alternatively, for miniaturized SPIONs enrichment, cells were scraped in 1 mL of 1X PBS and pelleted by centrifugation at 1,000 *g* for 5 min at 4 °C. Cell pellets were resuspended in 1mL of NP-40 buffer (0.1% NP-40, 200 mM HEPES/NaOH pH 7.4, 1X PIC), incubated on ice for 30 min and homogenized using a 2 mL dounce homogenizer (tight pestle) using 25 strokes. An aliquot was collected as the total fraction (TF). Next, nuclei and intact cells were pelleted by centrifugation at 500 *g* for 10 min at 4 °C, or 1000 *g* for 5 min at 4 °C (for miniaturized SPIONs), and the supernatant was transferred to a fresh tube. The pellet was resuspended in 4 mL or 1 mL (for miniaturized SPIONs) of isolation buffer, and cells were homogenized and centrifuged as mentioned above, respectively. The supernatants were pooled, resulting in the post-nuclear supernatant (PNS) fraction, which was used for lysosome enrichment using LS columns in combination with a QuadroMACS separator. The columns were equilibrated using 0.5% BSA in 1X PBS. Alternatively, for miniaturized SPIONS, µ columns placed on a µMACS separator, activated with 1 mL of degassed 70% ethanol and then equilibrated with 0.5% BSA in 1X PBS. The PNS was loaded onto the columns and the flow through (FT) was collected for subsequent analysis. Columns were washed with 3 mL of isolation buffer, the wash fractions were collected, columns were removed from the magnet, and lysosomes were eluted with 1 mL of isolation buffer using a plunger. Alternatively, lysosomes were eluted with 0.5 mL of elution buffer (1% Triton X-100, 40 mM HEPES/NaOH pH 7.4, 2.5 mM MgCl_2_, 10 mM Na_4_P_2_O_7_, 10 mM C_3_H_7_Na_2_O_6_P, and 1X PIC).

### Enrichment of Lysosomes by TMEM192-3xHA immunoprecipitation (TMEM IP)

HEK293 cells expressing TMEM192-3xHA were seeded in complete medium supplemented with 2 µg/mL puromycin. After 48 h, cells were washed twice with ice-cold 1X PBS, scraped in 1 mL of 1X PBS and centrifuged at 1,000 *g* for 5 min at 4 °C. The pellet was resuspended in KPBS buffer (136 mM KCl, 10 mM KH_2_PO_4_, pH 7.25 adjusted with KOH, 1X PIC) and homogenized using a 2 mL dounce homogenizer (tight pestle) using 25 strokes. Alternatively, for miniaturized TMEM IP, cells were harvested by trypsinization for 5 min at 37°C, trypsin inhibitor solution was added, detached cells transferred into a pre-cooled microtube and centrifuged at 1000 *g* for 5 minutes at 4 °C. The pellet was resuspended in NP-40 buffer, incubated on ice for 30 min and lyzed using a 2 mL dounce homogenizer (tight pestle) using 25 strokes. After the TF aliquot was collected, nuclei and intact cells were pelleted by centrifugation as mentioned above, and the PNS was transferred into a fresh tube containing 150 µL or 50 µL (for miniaturized TMEM IP) of pre-washed anti-HA magnetic beads. After incubation for 5 min at 4 °C in an end-over-end rotator, the tube was placed in a DynaMag Spin magnet, and the supernatant was collected. Beads were washed with KPBS buffer, and the last wash fraction was collected. Lysosomes were eluted from the beads using 50 µL of ice-cold elution buffer for 10 min on ice.

### β-Hexosaminidase Assay

Aliquots of individual lysosome enrichment fractions were loaded onto a 96-well plate in four replicates. To two wells, Triton X-100 (0.8% (v/v) final concentration) was added to disrupt the lysosomal membrane. Subsequently, reaction buffer (100 mM C_6_H_5_Na_3_O_7_, 10 mM para-nitrophenyl-*N*-acetyl-2-β-D-glucosaminide, 0.2% BSA, pH 4.6) was added to each well, followed by incubation for 15 min at 37 °C. The reaction was then stopped by addition of 0.4 M glycine pH 10.4 (final concentration), and absorbance was measured using a plate reader with excitation wavelength of 396 nm and emission wavelength of 405 nm.

### Immunoblotting

Samples were combined with Laemmli buffer^65^ (final concentration 1X), and incubated for 30 min at 45 °C. Proteins were separated by SDS-polyacrylamide gel electrophoresis (SDS-PAGE) with a constant voltage of 80 V for 2 h, and subsequently transferred onto a 0.45 µm nitrocellulose or PDVF membrane by wet blotting with a constant voltage of 70 V for 120 min. Membranes were blocked with 5% BSA in 1X Tris(hydroxymethyl)aminomethane-buffered saline containing 0.01% Tween-20 (TBS-T) for 1 h at room temperature (RT), and incubated with primary antibodies overnight at 4°C. The next day, membranes were washed three times with TBS-T, incubated with the respective secondary antibodies for 1 h at RT, and blots were imaged with the Enhanced chemiluminescence solution using a Fusion Solo 4M System.

### Sample Preparation for Mass Spectrometry

Sample protein concentration was determined using the DC protein assay. For lysosomes enriched by SPIONs/miniaturized SPIONs and TMEM IP/miniaturized TMEM IP, 50 µg and 10 µg were digested, respectively, using single-pot, solid-phase-enhanced sample preparation (SP3), as previously described^66^, with some modifications. Briefly, samples were reduced with 22 mM Dithiothreitol (DTT) for 30 min at 56 °C, and alkylated with 42 mM acrylamide for 30 min at RT, followed by quenching of the reaction with 22 mM DTT for 10 min at RT. Samples were then incubated with a 1:1 mixture of beads at a protein-to-bead ratio of 1:10 (w/w), followed by addition of 96% ethanol and incubation in a thermomixer at 1000 rpm for 5 min at 24 °C. Microtubes were placed on a magnetic stand, the supernatants were removed and beads were washed three times with 80% ethanol. Subsequently, samples were digested with sequencing-grade trypsin at an enzyme-to-protein ratio of 1:25 (w/w) in a thermomixer at 1000 rpm overnight at 37 °C. Tubes were transferred to a magnetic stand, the supernatant containing the peptides was transferred into a fresh tube, desalted using stop-and-go extraction (STAGE) tips as described elsewhere^67^, and dried using a vacuum centrifuge. For SPIONs-enriched samples, peptide concentration was determined using the quantitative fluorometric peptide assay.

### Liquid Chromatography-Tandem Mass Spectrometry Analysis

For SPIONs-enriched lysosomes, peptides were resuspended in 5% acetonitrile (ACN)/5% formic acid (FA) at a final concentration of 100 ng/µL, and 300 ng of each sample was analyzed. TMEM IP-enriched lysosomes were resuspended in 10 µL of 5% ACN/5% FA, and 30% of each sample was analyzed. For all LC-MS/MS analyses, a Dionex Ultimate 3000 nano-UHPLC coupled to an Orbitrap Fusion Lumos mass spectrometer was used operating in-house-produced analytical columns (45 cm spray tips generated from 360 µm outer diameter/100 µm inner diameter fused silica capillaries with a P-2000 laser puller and packed with with 3 µm ReprosiPur AQ C_18_ particles). Samples were loaded directly to the analytical column at a flow rate of 850 nL/min with 100% solvent A (0.1% FA), followed by separation with a 165 min linear gradient from 5% to 35% solvent B (95% ACN/0.1% FA). Eluting peptides were ionized in the positive mode and mass spectrometry (MS) analyses were performed in the data-independent-acquisition (DIA) mode. All scans were acquired in the Orbitrap analyzer. Following the acquisition of one MS1 scan, a total of 36 consecutive overlapping DIA MS2 scans were performed. For each scan, ions were isolated via the quadrupole with similar sized isolation windows (24.1 *m/z* each) covering the MS1 mass range from *m/z* 350 to 1,200. MS1 scans were acquired at a resolution of 120,000, a maximum injection time of 20 ms, and an automatic gain control (AGC) target of 5×10^5^, while MS2 scans were recorded at a resolution of 30,000, an *m/z* rage of 200 to 2,000, an AGC target of 1×10^6^ and a maximum injection time of 60 ms.

## QUANTIFICATION AND STATISTICAL ANALYSIS

### Mass Spectrometry Data Analysis

Thermo *.raw files were analyzed with Spectronaut 19.5.24 by direct DIA using UniProt Homo Sapiens (released 01/2024with 20,428 entries), in combination with the cRAP database containing common contaminants (https://www.thegpm.org/crap/). For each analysis, the following parameters were defined: enzyme: trypsin/P; number of missed cleavage sites: 2; minimum peptide length: 5 amino acids, mass tolerance: dynamic; fixed modification: propionamide at cysteine; variable modifications: oxidation at methionine and acetylation at protein N-terminus. The deep learning-assisted indexed retention time (iRT) regression^64^ was used with default settings for retention time alignment. Mass tolerances for matching of precursor/fragment ions and peak extraction windows were determined automatically by Spectronaut. Missing values were imputed within individual experiments (SPIONs, TMEM IPs), and results filtered with 1% false discovery rate (FDR) on the precursor and protein level (q-value < 0.01).

Spectronaut protein reports were analyzed using Python 3.12.8 using the NumPy^68^, pandas^69^, SciPy^70^, statsmodels^71^, matplotlib^72^ and seaborn^73^ libraries. For determination of numbers of identified total/lysosomal^18^ proteins, datasets without imputation were used, quantitative analyses were performed on data sets with imputed values. Furthermore, in order to enable lysosome-specific normalization in datasets strongly varying in complexity, additional replicate-wise normalization on the median intensity of v-ATPase subunits detected in both conditions with ≥ 10 unique peptides were performed. For further statistical analysis, normalized protein intensities were log_2_-transformed. Differential protein abundance between conditions (Miniaturized/Control) was assessed using a two-tailed Student’s t-test^74^ and the resulting p-values were adjusted for multiple testing using the Benjamini-Hochberg^75^ procedure. Proteins with log_2_ fold changes ≥ 1 and q-values < 0.05 were considered significantly differentially represented. Pearson correlation coefficients were calculated using v-ATPase-normalized protein intensity values. Data are plotted as mean ± SD, as mentioned in the figure legends. Samples sizes and replicate numbers are indicated as datapoints or mentioned in the figure legends.

### Data Processing and Statistical Analysis of β-Hexosaminidase Assays

HEXB assay results were analyzed using MS Excel 2019 and GraphPad Prism version 10.6. Data are plotted as mean ± SD as mentioned in the figure legends. The sample sizes are indicated as datapoints, and the number of replicates is indicated in the figure legends. Statistical analysis was performed using either Student’s t-test, one-way ANOVA, or two-way ANOVA. All statistical analyses were performed using GraphPad Prism. Statistical significance is denoted as * p < 0.05, ** p < 0.01, and *** p < 0.001 and **** = *p* < 0.0001.

## Figure legends

**Supplementary Figure 1:**
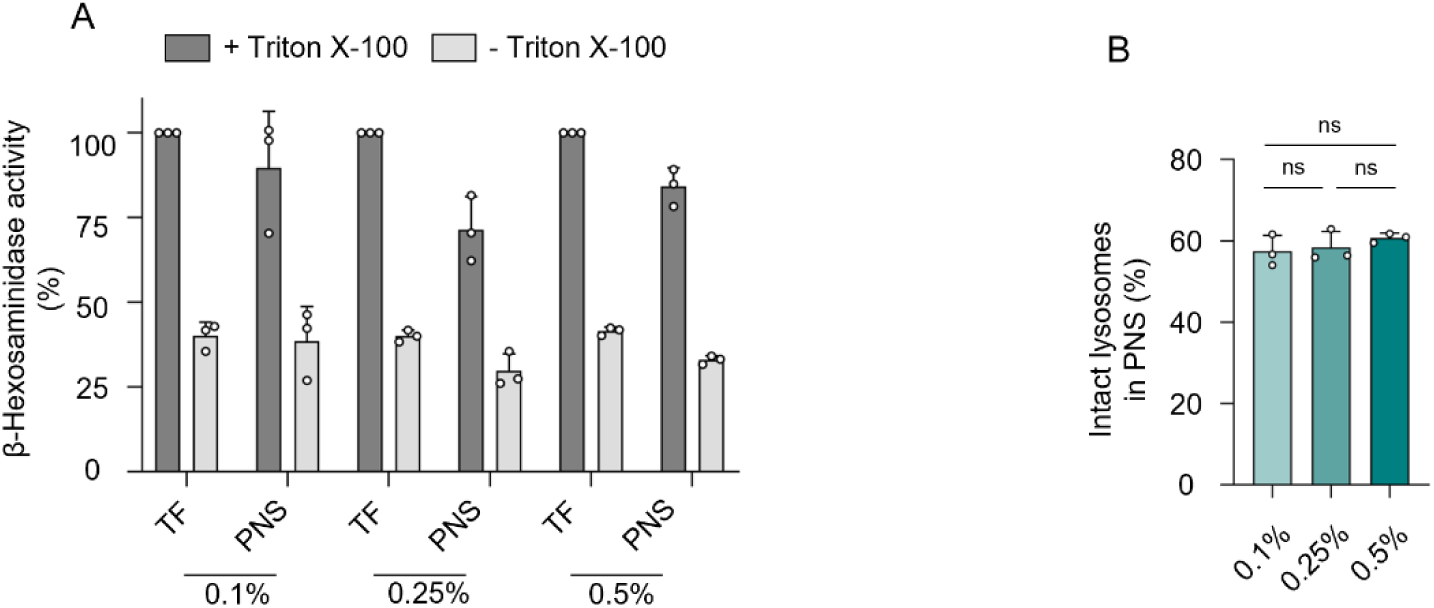
Release of intact lysosomes does not further improve with higher NP-40 concentration. (A/B) TF/PNS HEXB activity (A) and lysosomal intact ratio (B) from 6 million HEK293 cells incubated in 0.1%/0.25%/0.5% NP-40 prior to dounce homogenization. For raw values see Table S1. Shown are mean values ± SD, n = 3 (ns = not significant). TF = Total fraction, PNS = Post-nuclear supernatant.

**Supplementary Figure 2:**
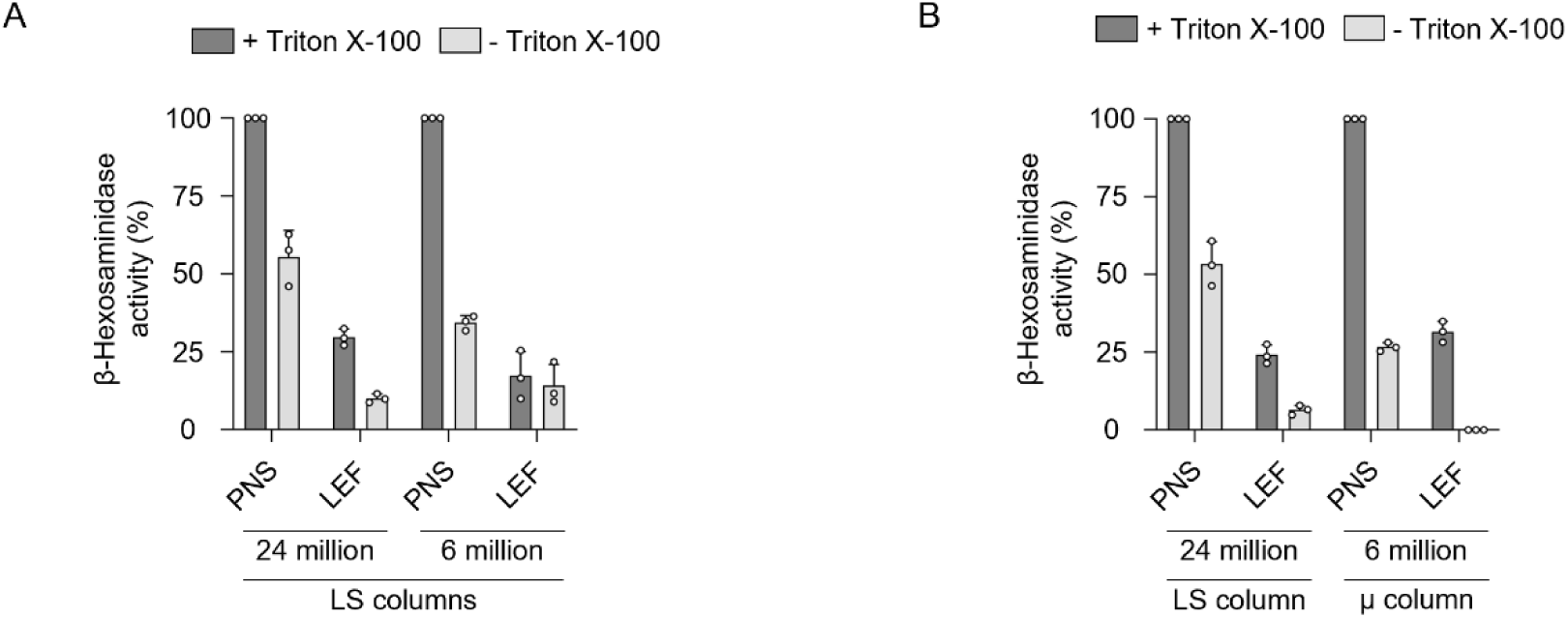
SPIONs enrichment of lysosomes from reduced cell numbers using plasma membrane destabilization. (A) PNS/LEF HEXB activity from SPIONs lysosome enriched of 24 million and 6 million HEK293 cells using LS columns. (B) PNS/LEF HEXB activity from 24 million cells enriched using LS columns and 6 million cells using µ columns. For raw values see Table S1. PNS = Post-nuclear supernatant, LEF = Lysosome-enriched fraction.

**Supplementary Figure 3:**
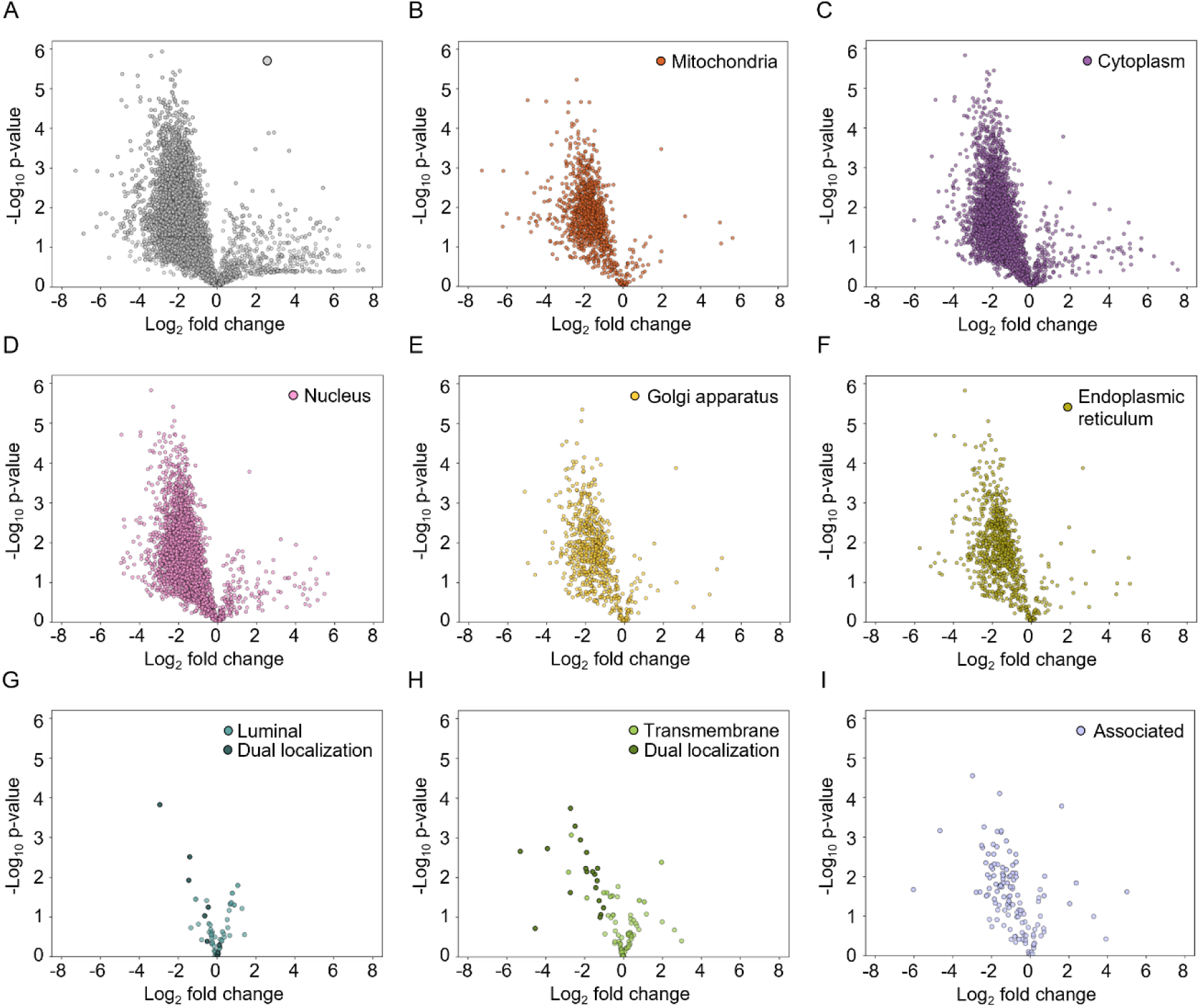
Mass spectrometry-based proteomics analysis of SPIONs-based lysosome enrichment from 24 million and 3 million cells. (A-I) Comparison of normalized relative protein abundances of SPIONs-based lysosome enrichment from 24 million cells and miniaturized SPIONs enrichment from 3 million cells. (A) total proteins detected. (B-F) proteins detected from distinct subcellular localizations: (B) mitochondria, (C) cytoplasm, (D) nucleus, (E) golgi apparatus, (F) endoplasmic reticulum. (G-I) Lysosomal proteins classified into (G) luminal, (H) transmembrane, and (I) associated proteins. Previously described proteins with dual localization to non-lysosomal compartments^31^ are highlighted as dark spots.

**Supplementary Figure 4:**
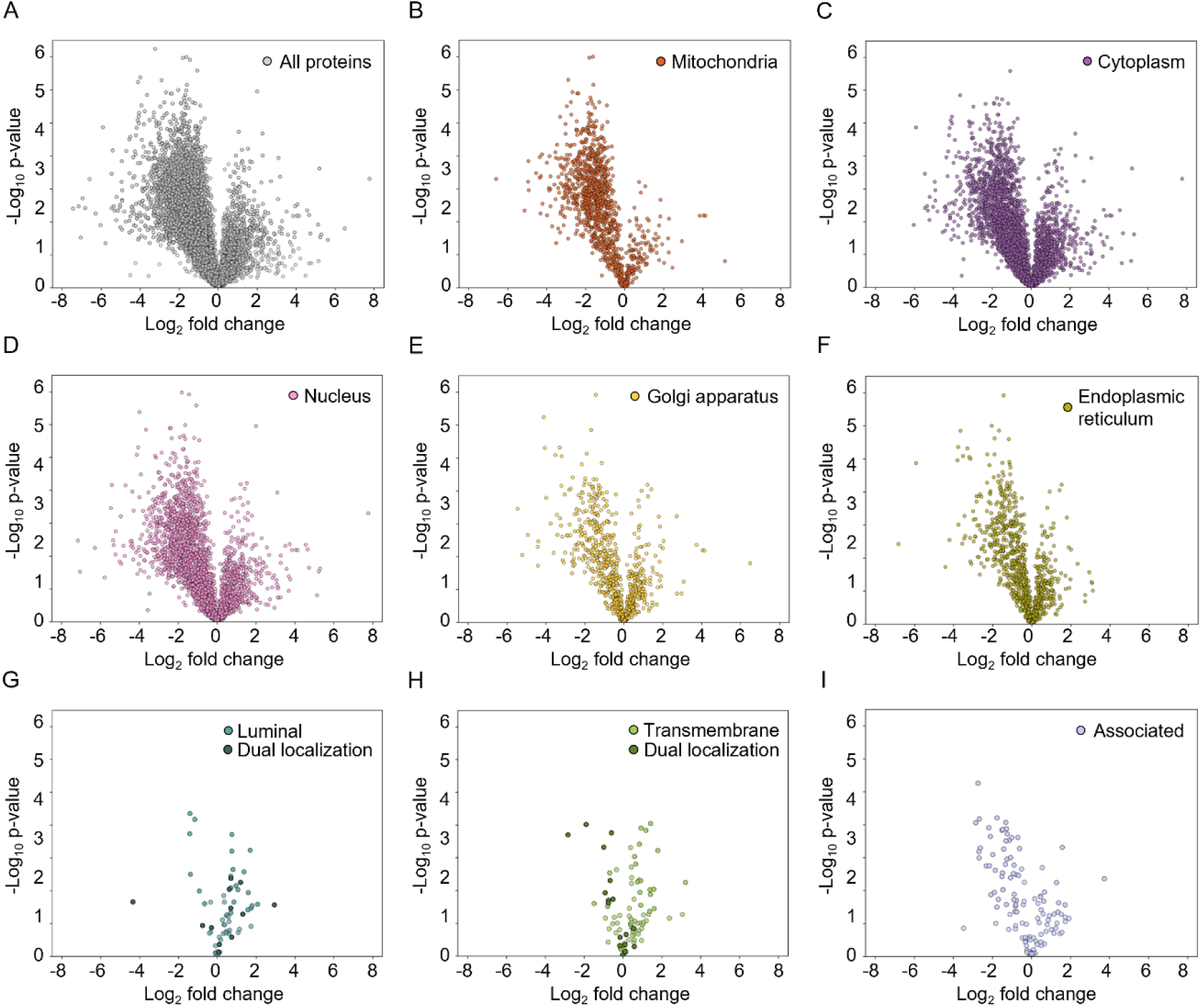
Mass spectrometry-based proteomics analysis of TMEM IP-based lysosome enrichment from 18 million and 0.5 million cells. (A-I) Comparison of normalized relative protein abundances of TMEM IP-based lysosome enrichment from 18 million cells and miniaturized TMEM IP enrichment from 0.5 million cells. (A) total proteins detected. (B-F) proteins detected from distinct subcellular localizations: (B) mitochondria, (C) cytoplasm, (D) nucleus, (E) golgi apparatus, (F) endoplasmic reticulum. (G-I) Lysosomal proteins classified into (G) luminal, (H) transmembrane, and (I) associated proteins. Previously described proteins with dual localization to non-lysosomal compartments^31^ are highlighted as dark spots.

## Supplementary Figure legends

### Table legends

**Supplementary Table 1:** β-Hexosaminidase assays results.

**Supplementary Table 2:** Protein identification from SPIONs lysosome enrichment experiments from 24 million and 3 million cells.

**Supplementary Table 3:** v-ATPase-normalized protein intensities from SPIONs lysosome enrichment experiments from 24 million and 3 million cells.

**Supplementary Table 4:** Distribution of non-lysosomal proteins in SPIONs lysosome enrichment experiments.

**Supplementary Table 5:** Protein identification from TMEM IP lysosome enrichment experiments using 18 million and 0.5 million cells.

**Supplementary Table 6:** v-ATPase-normalized protein intensities in TMEM IP lysosome enrichment experiments from 18 million and 0.5 million cells.

**Supplementary Table 7:** Distribution of non-lysosomal proteins in TMEM IP lysosome enrichment experiments.

